# *GhCO* and *GhCRY1* accelerate floral meristem initiation in response to increased blue light to shorten cotton breeding

**DOI:** 10.1101/2022.01.13.476244

**Authors:** Xiao Li, Yuanlong Wu, Zhenping Liu, Zhonghua Li, Huabin Chi, Pengcheng Wang, Feilin Yan, Yang Yang, Yuan Qin, Xuehan Tian, Hengling Wei, Aimin Wu, Hantao Wang, Xianlong Zhang, Shuxun Yu

## Abstract

The shoot apical meristem (SAM) is a special category of tissue with pluripotency that forms new organs and individuals, especially floral individuals. However, little is known about the fate of cotton SAMs as a tunica corpus structure. Here, we demonstrate that cotton SAM fate decisions depend on light signals and circadian rhythms, and the genes *GhFKF1*, *GhGI*, *GhCRY1* and *GhCO* were responsible for SAM fate decisions and highlighted via RNA sequencing (RNA-seq) analysis of different cotton cultivars, as confirmed by genetic analysis via the CRISPR–Cas9 system. *In situ* hybridization (ISH) analysis showed that the *GhCO* gene, induced by a relatively high blue light proportion, was highly upregulated during the initiation of floral meristems (FMs). Further blue light treatment analysis showed that the transition from vegetative to reproductive growth of SAM was promoted by a high proportion of blue light, coupled with high expression of the blue light-responsive genes *GhCO and GhCRY1*. Taken together, our study suggests that blue light signalling plays a key role in the fate decision of cotton SAM. These results provide a strategy to regulate the SAM differentiation of cotton by using the CRISPR–Cas9 system to change the ratio of red and blue light absorption to breed early-maturity cotton.

**One-sentence summary:** The SAM differentiation especially the initiation of floral meristem of upland cotton were mediated by genes *GhCO* and *GhCRY1* which in response to blue light.

## Introduction

Cells are the basic unit of life, and stem cells, are undifferentiated and possess remarkable pluripotency to replace damaged organs or form new organs and individuals throughout their lifespan both *in vivo* and *in vitro* (Tanaka et al., 2013). In flowering plants, most stem cells are apical meristem cells, including root and stem apical meristem (RAM and SAM) cells. At the postembryonic development stage, SAM and RAM cells generate aerial and underground parts, respectively (Aichinger et al., 2012; Kitagawa and Jackson, 2019). Stem cell differentiation, a physiological switch from a vegetative to reproductive stage, occurs early during the seedling phase is a multistep process driven by both intracellular signalling and extracellular cues (Liu et al., 2007; Turck et al., 2008; Srikanth and Schmid, 2011; Wang et al., 2018). Moreover, SAM stem cell fate determines floral meristem (FM) initiation and flowering (Jiang et al., 2013; Tanaka et al., 2013; Wagner, 2017; Kitagawa and Jackson, 2019).

Upland cotton is one of the most economically important crops worldwide. The early or late differentiation of FMs, which is determined by the fate of SAM, is directly related to the maturity period and architecture of a cultivar (Niwa et al., 2013), which is very important for practising double-cropping systems, namely, rapeseed/wheat-cotton production systems. One of the biggest challenges in cotton breeding is the long growth cycle (Li et al., 2021). Therefore, studying the fate determination mechanism of SAMs is very important to optimize the timing of cotton fruit branching and shorten the cotton growth stage to meet the large and increasing clothing demands of an ever-growing world population.

In *Arabidopsis*, previous studies showed that *CONSTANS* (*CO*) and *FLOWERING LOCUS T* (*FT*) were the central components for the transition from the vegetative to reproductive stage in flowering plants. *CO* is a zinc finger transcription factor (TF) that integrates flowering by activating *FT* expression in the afternoon under long days (LDs) (Song et al., 2015). FT, which acts as a long-distance signal in the light cycle, is a rapidly accelerated fibrosarcoma (RAF) kinase inhibitor-related protein that is transcribed in the leaves and then translates and migrates to the apical meristem through the vascular system after illumination (Corbesier et al., 2007; Turck et al., 2008). In the light spectrum, blue light is one of the most effective light signals in SAM differentiation, not only acting as a signal responding to stress (Lyu et al., 2021) but also participating in regulating many photomorphogenesis processes, including floral initiation and light entrainment of the circadian rhythm, hypocotyl elongation, seedling de-etiolation, stem elongation and leaf morphology (Thomas, 2006; Corbesier et al., 2007; Franklin, 2016).

Plants perceive light signals using various light receptors, including *KELCH REPEAT, F-BOX 1* (*FKF1*) (Liu et al., 2018), and cryptochrome (CRY) genes *CRY1* and *CRY2* (Ma et al., 2020; Wang and Lin, 2020). *FKF1* contains one N-terminal LOV domain, a target for ubiquitin-mediated degradation of the F box domain, and six protein–protein interaction tandem Kelch motifs (Nelson et al., 2000; Ito et al., 2012). The LOV domain, as the chromophore, comprised of flavin mononucleotide (FMN) binding sites, is responsible for light-induced protein–protein interactions between GIGANTEA (GI) and CO, which regulate the expression of *CO* and *FT* and affect flowering time in response to photoperiod and circadian clock signals (Thomas, 2006; Ito et al., 2012). FKF1 also binds to proteolytic targets, a Dof TF, *CYCLING DOF FACTOR 1* (*CDF1*), which binds to the *CO* promoter, inhibiting the transcription of *CO* (Song et al., 2015). Additionally, highly conserved flavoprotein CRYs, comprising a conserved N-terminal photolyase homologous region (PHR) domain and an unstructured cryptochrome C-terminal extension (CCE) effector domain (Ma et al., 2020; Wang and Lin, 2020), are degraded by light-dependent ubiquitination (Liu et al., 2016). *CRYs* have been investigated to facilitate *FT* expression in response to blue light by suppressing degradation of the CO protein (Zuo et al., 2011).

However, in comparison with extensive studies in the model plant *Arabidopsis*, the physiological impact and mechanisms of light in the maintenance of SAM pluripotency in feed and cash crops remain to be investigated. Here, the gene response to light signals was highlighted during the transition from vegetative to reproductive growth of cotton SAMs using RNA-seq and qRT–PCR. Additionally, we confirmed the roles of *GhCRY1* and *GhCO* in mediating SAM fate decisions, especially those of FMs, and revealed that *GhCRY1* and *GhCO* were regulated by blue light and that *GhCRY1* accelerated flowering. In sharp contrast, *GhCO* played a role in balancing the number and fate of floral primordium differentiation at meristems, different from *Arabidopsis*, which acted in the phloem (An et al., 2004; Turck et al., 2008). Notably, we further investigated the influences of different red:blue (R:B) light ratio treatments on cotton SAM differentiation and flowering, showing that a high proportion of blue light evokes early flowering of cotton, suggesting a critical role of blue light in adjusting floral and fruit branch primordium initiation. Notably, agronomic trait assays in the spectral incubator demonstrated a feasible way to breed early-flowering cotton cultivars by remodelling the blue light signalling activities mediated by *GhCRY1* and *GhCO*.

## Results

### Dynamics of cotton SAM differentiation revealed by anatomical analysis

The growth and development process of cotton are dependent on the division and differentiation of the stem cell population in the SAM, which is generally divided into three stages: the vegetative stage, SAM initiation stage and reproductive stage (Fig. 1A). Briefly, the fate of cotton SAM cells, including those in the leaf primordium (LP), branch meristem (BM) and FM, determined the flowering time and plant architecture (Fig. 1A) (Bhalla and Singh, 2006; Aichinger et al., 2012).

**Fig. 1.**
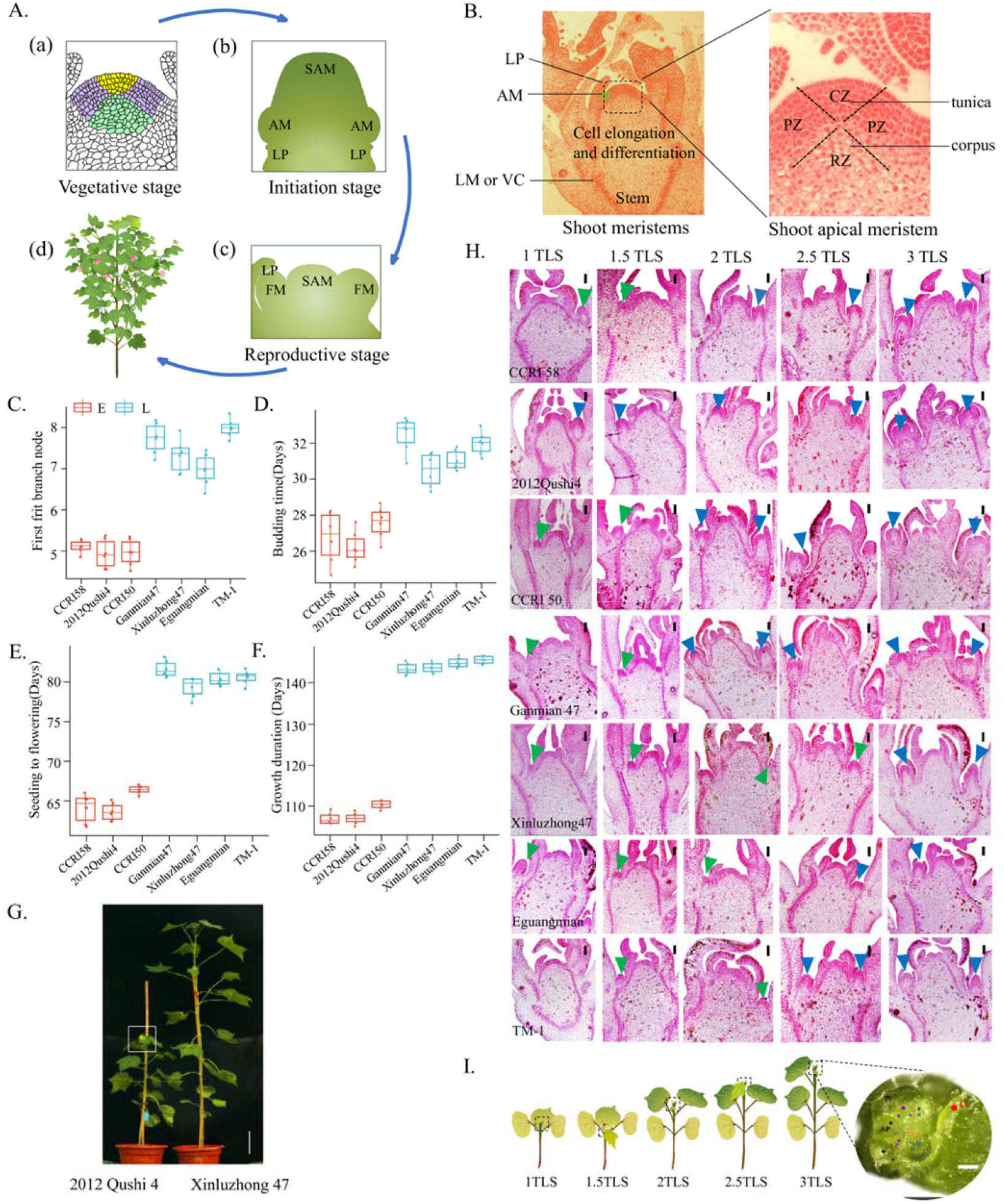
Dynamic changes in cotton shoot apical meristem (SAM) differentiation. **(A)** Schematic diagram of cotton growth and development stages. (a) Model of cotton stem apical meristem during the vegetative stage. Yellow area, central zone (CZ); green area, rib zone (RZ); purple area, peripheral zone (PZ). (b) The transitional stage of vegetative to reproductive growth, which is the initial period of fate determination in the cotton apical meristem. SAM, shoot apical meristem; AM, axillary meristem; LP, leaf primordium. (c) Model of the apical meristems during the reproductive stage in which the SAM fate was decided, and the axillary meristem differentiated into floral meristems (FMs). (d) Cotton that has blossomed and borne fruit. **(B)** The tunica and corpus structure of cotton SAMs. The left panel shows paraffin sections of cotton shoot meristems. LM, lateral meristem; VC, vascular cambium. The right panel represents cotton SAMs. **(C-F)** Phenotypic analysis of three early- and four late-maturity cotton cultivars at the first fruit branch node (C), the budding time (D), the days from seedling to flowering (E) and the days of whole growth periods (F). E, early-maturity variety; L, late-maturity variety. **(G)** The plants were derived from cotton cultivars of different maturities. The left panel shows the early-maturity cultivar Q, and the right panel shows the late-maturity cultivar XLZ. Scale bars, 10 cm. **(H)** The dynamics of cotton SAM differentiation, where 1, 1.5, 2, 2.5 and 3 indicate the first true leaf stage (TLS), stage between the first and second TLS, the second TLS, stage between the second and third TLS and the third TLS, respectively. Yellow triangle, vegetative primordium; blue triangle, reproductive primordium; scale bar, 100 μm. **(I)** Experimental scheme of RNA-seq experiments. Seedlings of early- and late-maturity cotton were grown under LD conditions (16 h light/8 h dark) at 28 °C. The dotted box indicates the position of SAMs collected for the mRNA-seq experiment. The plot on the right is a magnified apical meristem of cotton under a stereomicroscope. Red oval, LP; blue oval, AM; black oval, sepal primordium (SP); purple oval, FM; orange oval, SAM. Scale bar, 300 μm.

To determine the anatomical structure of SAMs, cross sections were obtained by sampling the stem shoots from vegetative to reproductive stages, and it was found that SAM stem cells were a typical tunica-corpus structure (Fig. 1B). To investigate the mechanism of cotton SAM differentiation, the phenotypes of three early-maturing cotton cultivars, namely, CCRI58 (C), 2012Qushi4 (QS) and CCRI50 (CR), and four late-maturing cultivars, Ganmian47 (G), Xinluzhong47 (XLZ), Eguangmian (EG) and TM-1 (TM), were investigated and showed that the first fruit branch node, budding time, flowering time, whole growth period, and height of plants were significantly different between early-maturing and late-maturing cultivars (Fig. 1C-G and Table S1). Showing that the early-maturing cultivars budding, flowering and maturing much earlier than the late-maturing cultivars.

To further investigate the SAM dynamics, five developmental stages, including the first true leaf stage (1 TLS), the stage between the first and the second true leaf stage (1.5 TLS), the second true leaf stage (2 TLS), the stage between the second and the third true leaf stage (2.5 TLS), and the third true leaf stage (3 TLS), were sampled and compared (Fig. 1H-I). Morphological analysis showed that the differentiation of the FM varied between early- and late-maturity cultivars. Briefly, the FM differentiated earlier in QS than in XLZ; the former occurred at 1 TLS, and the latter occurred at 3 TLS (Fig. 1H). The other two early-maturity cultivars, C and CR, differentiated at 2 TLS, and the two late-maturity cultivars EG and TM differentiated at 2.5 TLS (Fig. 1H). These results showed that during the transition of SAM (initiation) from vegetative growth to reproductive development, the floral primordia differentiation of early-maturing cultivars occurred much earlier than that of late-maturing cultivars, suggesting that the spatiotemporal differentiation of SAM cells determines cotton flowering time.

### Light and circadian rhythms play an important role in fate decisions in cotton SAMs during the transition from vegetative to reproductive growth

To explore the molecular mechanism of SAM development in cotton, shoot tips were collected at five different developmental stages to identify differentially expressed genes (DEGs) between seven different cotton varieties by using RNA sequencing (RNA-seq) (Fig. 1I and Fig. S1). The expression levels of genes were determined by calculating the number of unique matched reads for each gene and then normalizing this number to fragments per kilobase of transcript per million mapped reads (FPKM), and a total of 47303 genes were expressed in at least one sample during cotton SAM development (FPKM ≥ 1) (Mortazavi et al., 2008). Distribution analysis of all genes showed that they were distributed equally among the At and Dt genomes (At: 34379 genes, 49.34%; Dt: 35298 genes, 50.66%), and the same trend was found for the 47303 expressed genes (At: 23189, 49.02%; Dt: 24114, 50.98%) (Fig. 2A). However, there was a slightly lower proportion of genes with low expression (FPKM < 1) at 1.5 TLS in the early-maturity sample and 2.5 TLS in the late-maturity sample compared to other four stages among At and Dt genomes, suggesting that gene expression was promoted at 1.5 TLS in the early-maturity cultivars and 2.5 TLS in the late-maturity cultivars (Fig. 2B). The Pearson correlation coefficients (PCCs) of gene expression (FPKM) between stages of different cotton cultivars were highly correlated (cor > 0.6, p < 0.0001, Fig. 2C-D, Table S2 and S3). These results showed that there were no subgenome differences in the expression genes during cotton SAM development.

**Fig. 2.**
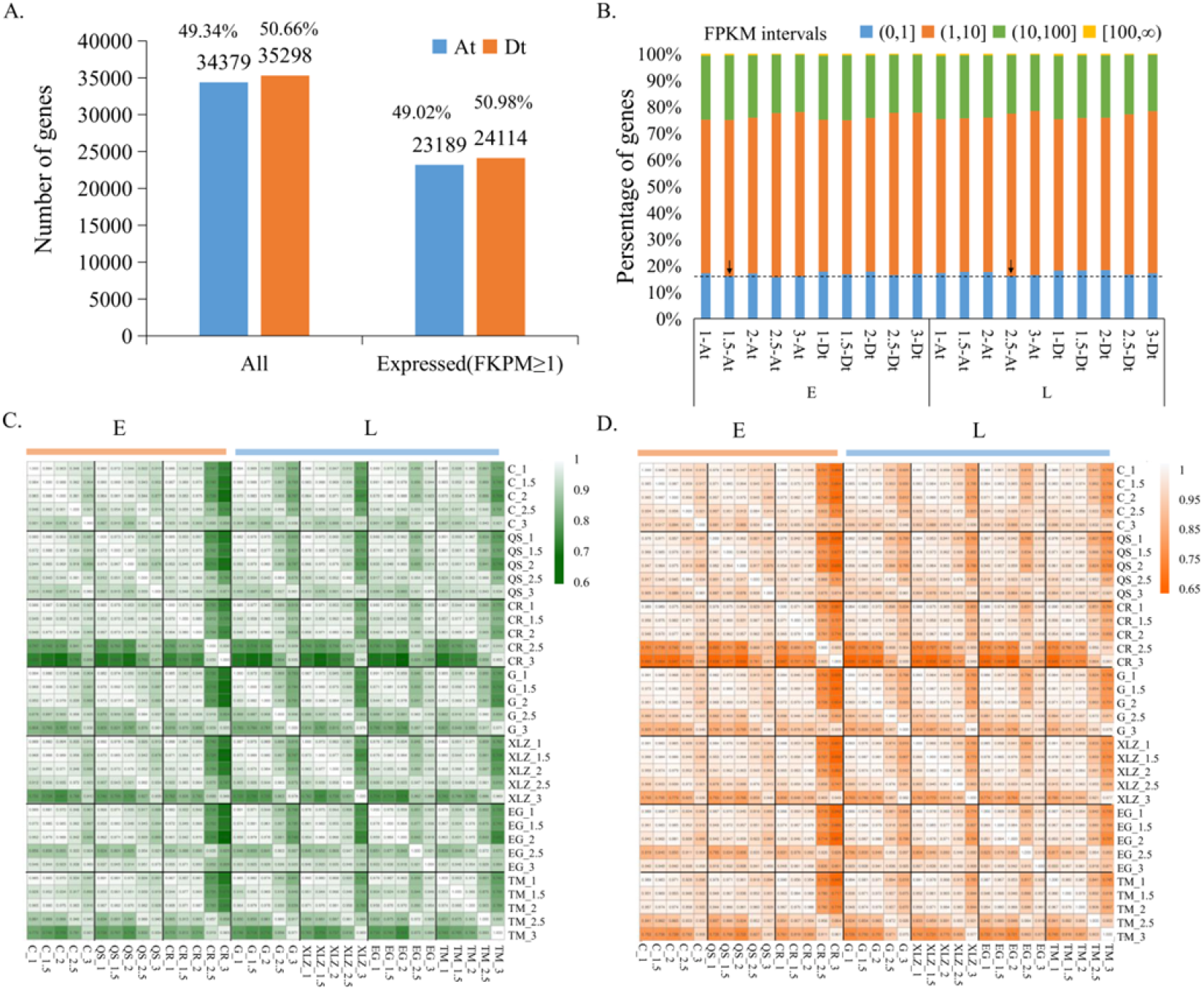
Overview of the cotton shoot apical meristem gene expression profiles at five developmental stages. **(A)** Total number of annotated genes in the At (left) and Dt (right) subgenomes and the number of genes that were expressed in at least one sample (FPKM ≥ 1). **(B)** The proportion of genes with the indicated expression strength at each developmental stage. The strength of expression is divided into four categories according to the normalized expression level (fragments per kilobase of exon model per million mapped reads, FPKM). **(C-D)** The Pearson correlation coefficients (PCCs) of gene expression (FPKM) between stages. The PCCs of expressed genes from the At (C) and Dt (D) subgenomes were calculated separately. 1, 1.5, 2, 2.5 and 3 TLS indicate the first leaf stage, the stage between the first and second true leaves, the second true leaf stage, the stage between the second and third true leaf stages and the third true leaf stage, respectively. TLS: true leaf stage; E, early-maturity variety; L, late-maturity variety.

Then, principal component analysis (PCA) was performed on all 35 samples and found that the high-density time series transcripts could be divided into five categories according to the developmental stages, although the 1.5 and 2 TLS groups highly overlapped (Fig. 3A-B), implying that there were differences in different developmental stages. To fully understand the DEGs, 4561 DEGs were identified by filtering with edgeR (P < 0.05 and log2 |fold-change| > 2 in normalized expression levels) (Fig. 3C-D), showing that the number of DEGs in different varieties at the same developmental stages was low, but in different developmental stages of the same cultivar, the number of DEGs was large. Moreover, comparison between the early- and late-maturity cultivars showed that there were 2408 overlapping DEGs (Fig. 3E), which were significantly involved in ‘Response to karrikin’, ‘Response to light stimulus’, ‘Response to circadian clock and flowering’, ‘Response to far red light and blue light’ and ‘Response to hormone’ Gene Ontology (GO) terms (Fig. 3F). The expression trend analysis showed that two gene expression patterns (profiles 8 and 9) were significantly enriched among the 4561 DEGs (Fig. S2A). Expression level analysis showed that in profile 8, the level increased gradually with development, but the change was inverse in profile 9 (Fig. S2B-C). Consistent with the above GO enrichment results, DEGs involved in profiles 8 and 9 were enriched in the same pathways (Fig. S2B-C). To further verify whether light and circadian rhythms regulate FM differentiation time, two genes (*GhGI* and *GhFKF1*) involved in these pathways were identified, and the expression patterns of these genes were detected in leaves and stems at different times (0:00, 6:00, 9:00, 12:00, 15:00, 18:00, 21:00, 24:00) of the day and showed that these genes were responsive to circadian rhythms (Fig. 3G-H). Suggesting that the transition from vegetative to reproductive growth in response to light signal and circadian rhythm.

**Fig. 3.**
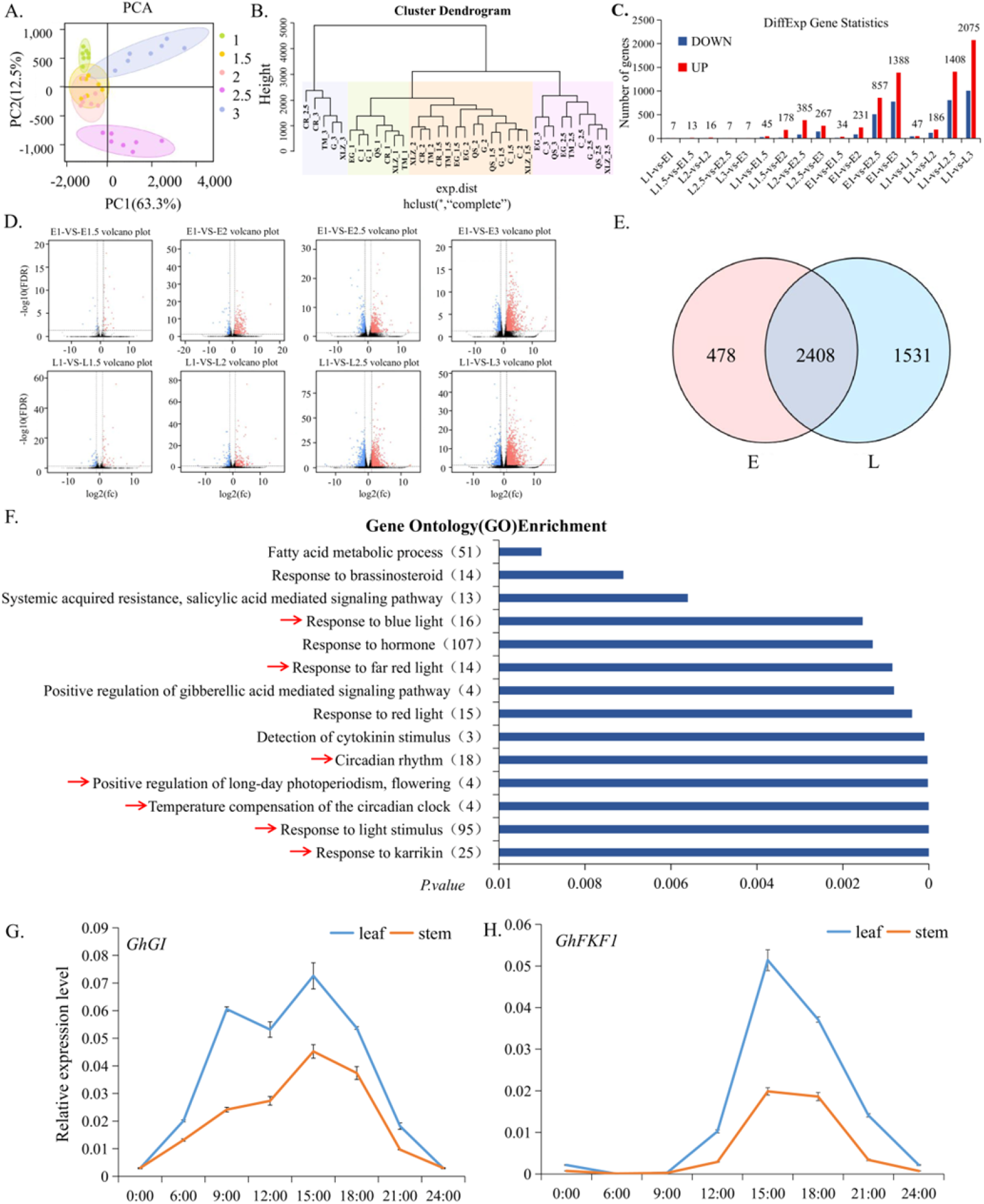
Transcriptome profiling of cotton SAM cell clusters. **(A)** Principal component analysis (PCA) of RNA-Seq data. **(B)** The clustering tree diagram shows different clustering groups. **(C)** The numbers of up- and downregulated differentially expressed genes (DEGs) for each pairwise comparison. **(D)** Volcano plots to visualize DEGs in early- and late-maturity cotton seedlings by comparing the same growth stage between different maturities and the same maturity between different growth stages. Red dots indicate significantly upregulated genes (log2(fold change) ≥ 0.75 and p < 0.05). Blue dots indicate significantly downregulated genes (log2(fold change) ≤ −0.75 and p < 0.05). **(E)** A Venn diagram shows the overlapping number of DEGs between the indicated samples. **(F)** Significant GO term enrichments for the 2408 overlapping DEGs. The number of genes in each GO term is indicated in the brackets. The blue arrows represent key metabolic pathways in which the genes *GhGI* and *GhFKF1* were involved. **(G-H)** Relative expression level analysis of leaves and stalks separately at 2 TLS at different times (0:00, 6:00, 9:00, 12:00, 15:00, 18:00, 21:00, 24:00) of the day by qRT–PCR of the genes *GhGI* (G) and *GhFKF1* (H), which are involved in responding to circadian rhythm and flowering. The plants were grown under LD conditions (16 h light/8 h dark). The error bars represent standard deviations (SD, n ≥ 3). TLS, true leaf stage; E, early-maturity variety; L, late-maturity variety.

Furthermore, weight gene co-expression network analysis (WGCNA) was performed on all 4156 DEGs and showed that there were 5 different modules, corresponding to the 5 co-expression networks (Fig. 4A). There was a significantly negative correlation between the turquoise and brown modules (Fig. 4B). Interestingly, four of the flower-related genes identified by genome-wide association analysis of 355 group cultivars (Li et al., 2021) were differentially expressed between early- and late-maturity cultivars during cotton SAM development, and the expression level peak occurred at 2 or 2.5 TLS (Fig. 4C). One of the genes (*GhELF4*) was involved in the turquoise module and co-expressed with *GhGI*, *GhFKF1* and other genes (Fig. 4D). Additionally, rhythm expression level analysis of other co-expressed genes in the turquoise module, such as *GhAGL24*, *GhSOC1*, *GhELF3*, *GhPHOT2*, and *GhVIP5*, in leaves and stems showed that all of these genes respond to circadian rhythms (Fig. 4E-P). These results suggest that *GhGI* and *GhFKF1* may participate in the circadian rhythm regulation of cotton flowering.

**Fig. 4.**
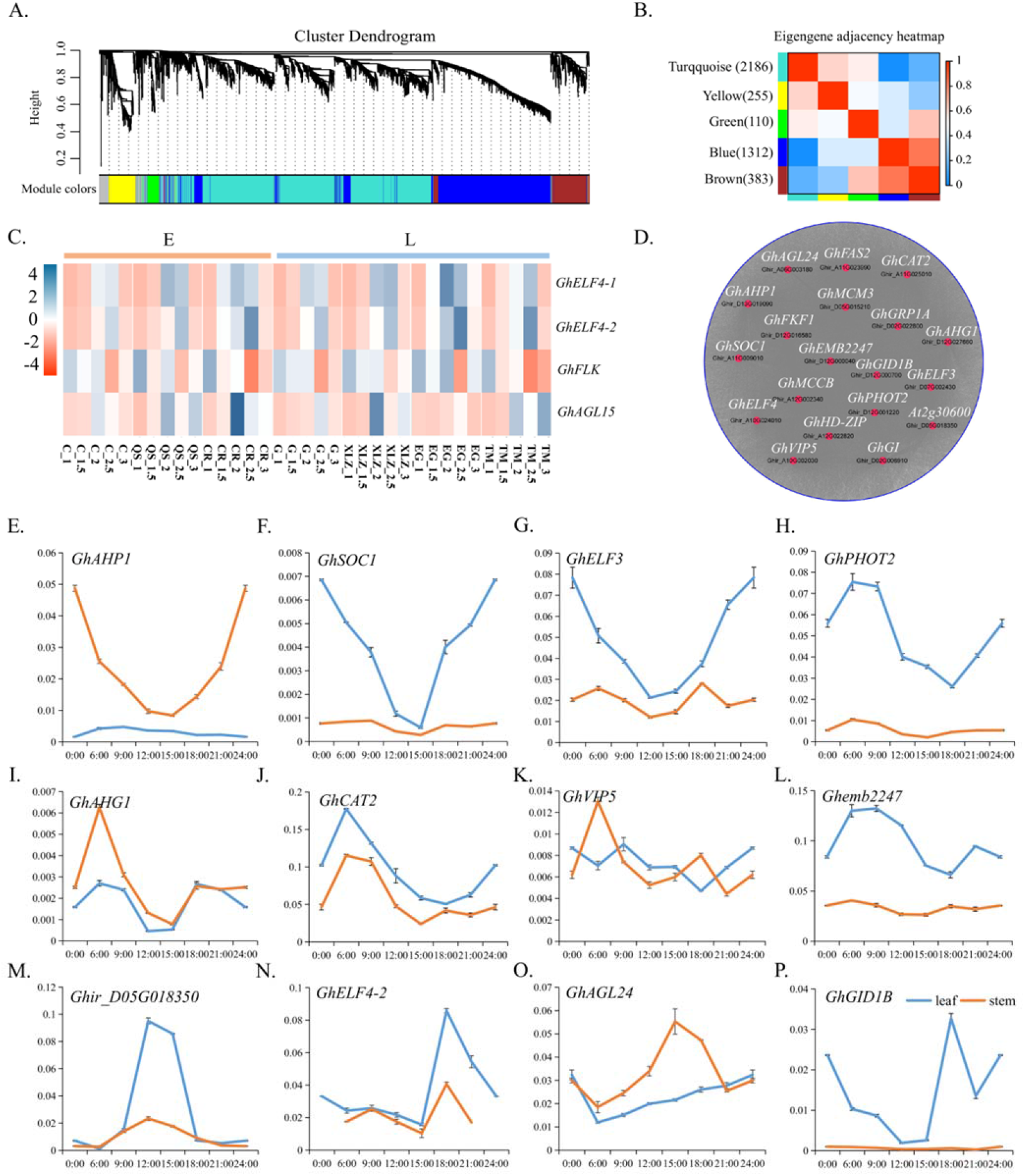
Floral meristem initiation of cotton in response to rhythm and circadian clock. **(A)** Hierarchical cluster tree showing co-expression modules identified by WGCNA. Each leaf in the trees is one gene. The major tree branches constitute 5 modules labelled by different colours. **(B)** Module-sample association. Each row corresponds to an intersection that indicates the correlation coefficient between the module and sample. **(C)** Expression level analysis of the genes *GhELF4-1*, *GhELF4-2*, *GhFLK* and *GhAGL15* belonging to the overlapping genes between our DEGs and previous genome-wide association analysis results (Li et al., 2021). E, early-maturity variety; L, late-maturity variety. **(D)** WGCNA for turquoise co-expression modules by Cytoscape showing a significant enrichment of known boundary-specific genes in response to flowering development. Red indicates reproductive development-related regulators defined according to the GO notion. Dots around the circle represent other co-expressed genes. Lines represent relationships. The white and black font represent the gene abbreviation and ID, respectively. **(E-P)** Relative expression level analysis by qRT–PCR of genes that co-expressed not only *GhGI* but also *GhELF4* in leaves and stalks at 2 TLS at different times (0:00, 6:00, 9:00, 12:00, 15:00, 18:00, 21:00, 24:00) of the day. The samples were grown under LD conditions (16 h light/8 h dark). Data are represented as the mean ± SD (n ≥ 3). TLS, true leaf stage.

To confirm this hypothesis, mutants of *Ghfkf1* and *Ghgi* were created by CRISPR–Cas9 with two small guide RNAs (sgRNAs) (Fig. 5A-C), and the mutants were determined by the Hi-TOM platform (Liu et al., 2019) to track mutations (Fig. 5D-E). Phenotypic analysis of the mutants showed that there was no lateral branch differentiation in the *Ghfkf1* mutant, but in the *Ghgi* mutant, more branches and flower buds were found (Fig. 5F-G). This result implies that during cotton SAM initiation from vegetative to reproductive stages, the *GhFKF1* gene promoted lateral differentiation, but *GhGI* inhibited floral primordia differentiation, possibly related to light stimulus and circadian rhythm.

**Fig. 5.**
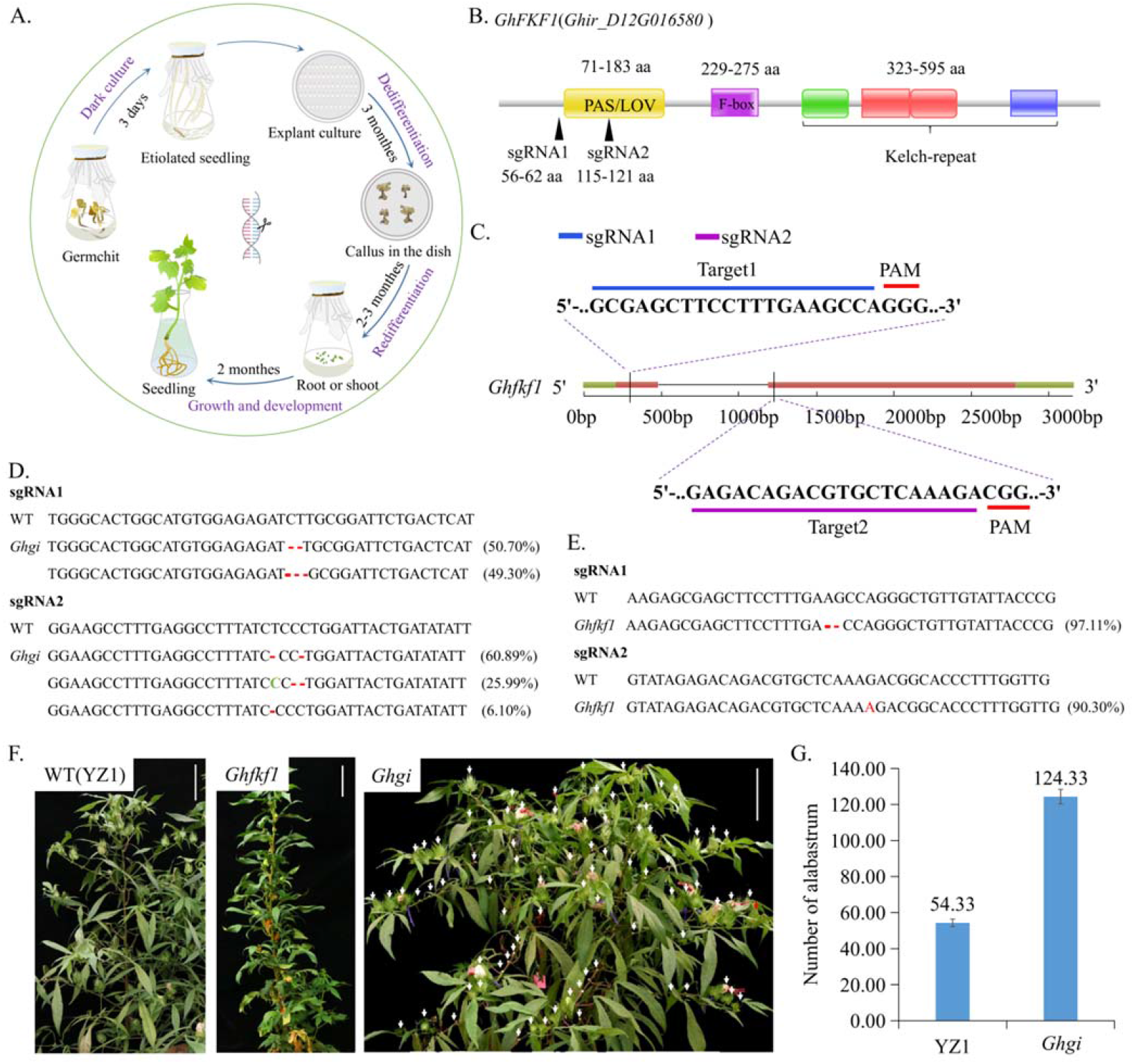
*Ghfkf1* and *Ghgi* mutants created by the CRISPR–Cas9 strategy. **(A)** Diagram showing the process of *Ghco* mutant plant production by the CRISPR–Cas9 strategy. **(B)** Schematic of the two small guide RNAs (sgRNAs, blue and purple bars) designed to target *GhFKF1* loci. Protospacer adjacent motifs (PAMs) are indicated by red bars. **(C)** Schematic of the two small guide RNAs (sgRNAs, blue and purple bars) designed to target *GhFKF1* loci. PAMs are indicated by red bars. **(D-E)** Sequencing results showing indel mutations in the *GhGI* (D) and *GhFKF1* (E) knockout cotton plants. The percentages in parentheses represent gene editing efficiency on the right of sequences. Red bars represent missing sites, and the green SNPs represent the mutation sites. **(F)** Phenotypes of *Ghfkf1* and *Ghgi* edited plants. The white arrows indicate flowers or flower buds. Scale bar, 10 cm. **(G)** The statistics of the alabastrum numbers of WT plants YZ1 and edited plants *Ghgi*. Data are represented as the mean ± SD (n = 3).

### *GhCO* responds to blue light and transits SAM stem cell differentiation into the floral primordium in rate and number

A previous study showed that many TFs were involved in SAM fate decisions, especially in FM determinacy, by responding to light (Komeda, 2004; Liu et al., 2007; Turck et al., 2008; Song et al., 2015; Lai et al., 2021). To better understand the function of light signals in SAM differentiation during the transition from vegetative to reproductive stages, a total of 405 TFs were identified from the 4561 DEGs according to their functions and divided into four groups according to changes in expression (Fig. S3A and Table S4). Abundance bubble diagram analysis showed that TF families such as *ERF*, *MYB*, *C2H2*, *HD*-*ZIP*, *C3H*, *TALE*, *MYB*, *NAC* and *CO*-*like* were enriched (Fig. S3B), among which the *HD*-*ZIP*, *CO*-*like*, *TALE* and *RAV* families playing important roles in flowering were identified (Fig. 6A and Table S5).

**Fig. 6.**
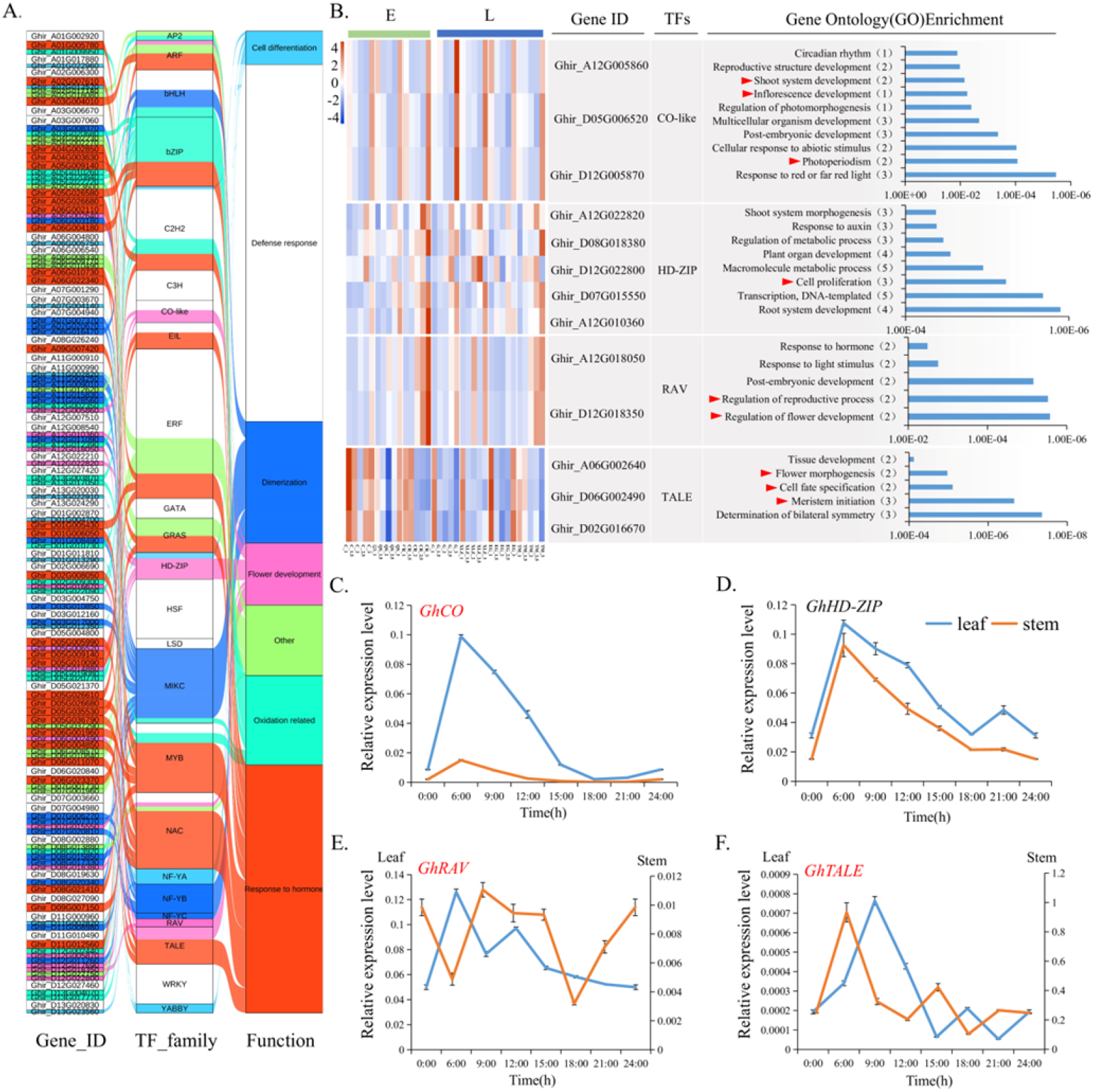
The transcription factors (TFs) analysis of DEGs. **(A)** Sankey diagram of the exact functional annotation of TFs. **(B)** Heatmap of 13 TFs associated with floral development. Related TFs and significant GO enrichments (right) are presented. **(C-F)** Relative expression level analysis by qRT–PCR of TFs in response to flower development GO terms at leaves and stalks during 2 TLS at different times (0:00, 6:00, 9:00, 12:00, 15:00, 18:00, 21:00, 24:00) of the day. The samples were grown under LD conditions (16 h light/8 h dark). Data are represented as the mean ± SD (n ≥ 3). TLS, true leaf stage; E, early-maturity variety; L, late-maturity variety.

Moreover, the expression and function analysis showed that *CO*-*like* TFs were significantly enriched in ‘response to red or far red’, ‘photoperiodism’,‘postembryonic development’, ‘inflorescence development’, ‘shoot system development’ and ‘reproductive structure development’, with a peak expression level at 3 TLS. *GhHD*-*ZIP*, which was mainly enriched in ‘root system development’ and ‘cell proliferation’, was upregulated during SAM development. The *GhRAV* involved in ‘regulation of flower development’ and ‘regulation of reproductive process’ reached a peak in expression at 3 TLS. *GhTALE*, which was mainly enriched in ‘determination of bilateral symmetry’, ‘meristem initiation’, ‘cell fate specification’ and ‘flower morphogenesis’, was downregulated during SAM development, peaking at 1 TLS (Fig. 6B). Interestingly, the rhythm expression analysis of these four TFs showed that they respond to circadian rhythm, and *GhCO*-*like*, *GhRAV* and *GhTALE* were preferentially expressed in leaves (Fig. 6C-F). These results indicated that the TFs *GhCO*-*like*, *GhRAV* and *GhTALE*, related to circadian rhythm regulation and light signals, may be involved in SAM development.

To further clarify the function of light signalling during cotton SAM development, the expression patterns of several flowering-related genes involved in the light pathway were detected qRT–PCR (Fig. S3A-B) (Nelson et al., 2000; Zuo et al., 2011; Wang and Lin, 2020). The results corresponded well to the expression level derived from the RNA-seq data (Fig. S3C).

To demonstrate the light signalling function in the SAM stem cell fate decision and differentiation of the FM, *in situ* hybridization (ISH) was performed and showed that *GhCO*-*like* was expressed in the SAM, LP and lateral meristem cells (Fig. 7A-B). During the development of SAMs, at the reproductive stage, the expression of *CO*-*like* was much stronger than that at the vegetative stage and the post-reproductive stage, coincidental with the transcriptome analysis result that the expression level reached a peak at 3 TLS when the reproductive stage was initiated (Fig. 6B and 7A-B). The results suggested that *GhCO*-*like* may function in promoting SAM initiation from the vegetative stage to the reproductive development stage. To further verify how *GhCO*-*like* responds to light signals to affect the differentiation fate of SAMs, different R:B ratio treatments were performed, and it was found that under different R:B ratio treatments, the higher the blue light proportion was, the stronger the *GhCO* signal accumulation in the SAM stem cells (Fig. 8A-B). These results indicate that the expression of *GhCO* was induced by the blue light signal in SAM stem cells, which may promote the transition from vegetative to reproductive growth of the SAMs.

**Fig. 7.**
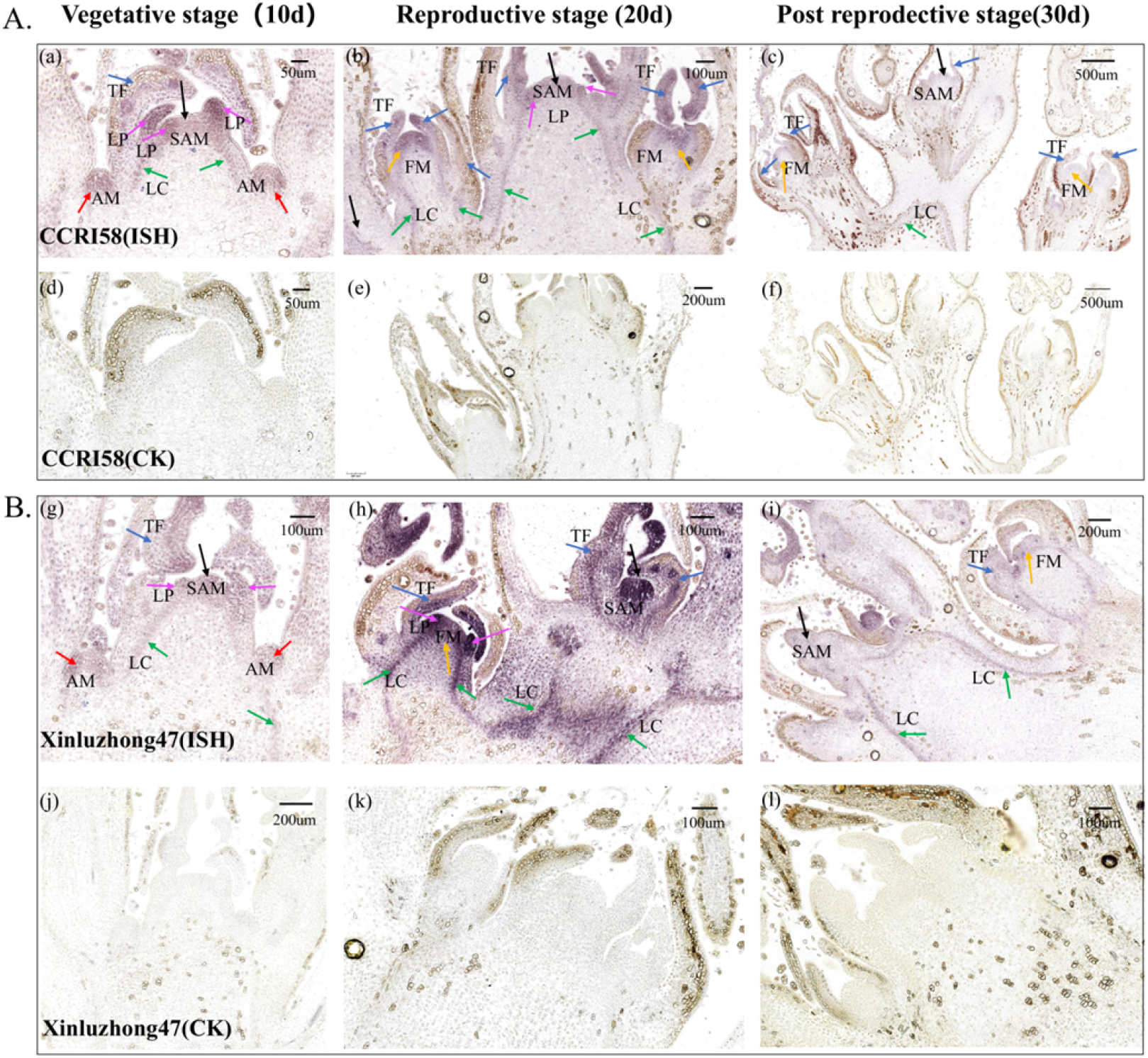
*In situ* hybridization (ISH) was used to localize *GhCO* in the developing SAMs of cotton. Amaranth staining indicates the presence of ISH-amplified target gene transcripts. **(A)** The localization of *GhCO* in the developing SAMs of cotton maturity C. **(B)** The localization of *GhCO* in the developing SAMs of cotton maturity XLZ. (a-c and g-i) Representative micrographs of longitudinal sections of SAMs for *GhCO*, with constitutive expression in the leaf primordium (LP, the pink arrow), axillary meristem (AM, the red arrow), floral meristem (FM, the orange arrow), shoot apical meristem (SAM, the black arrow) and tender leaf (TL, the green arrow). (d-f and j-l) Negative controls of C (d-f) and XLZ (j-l) omitting the reverse transcription step.

**Fig. 8.**
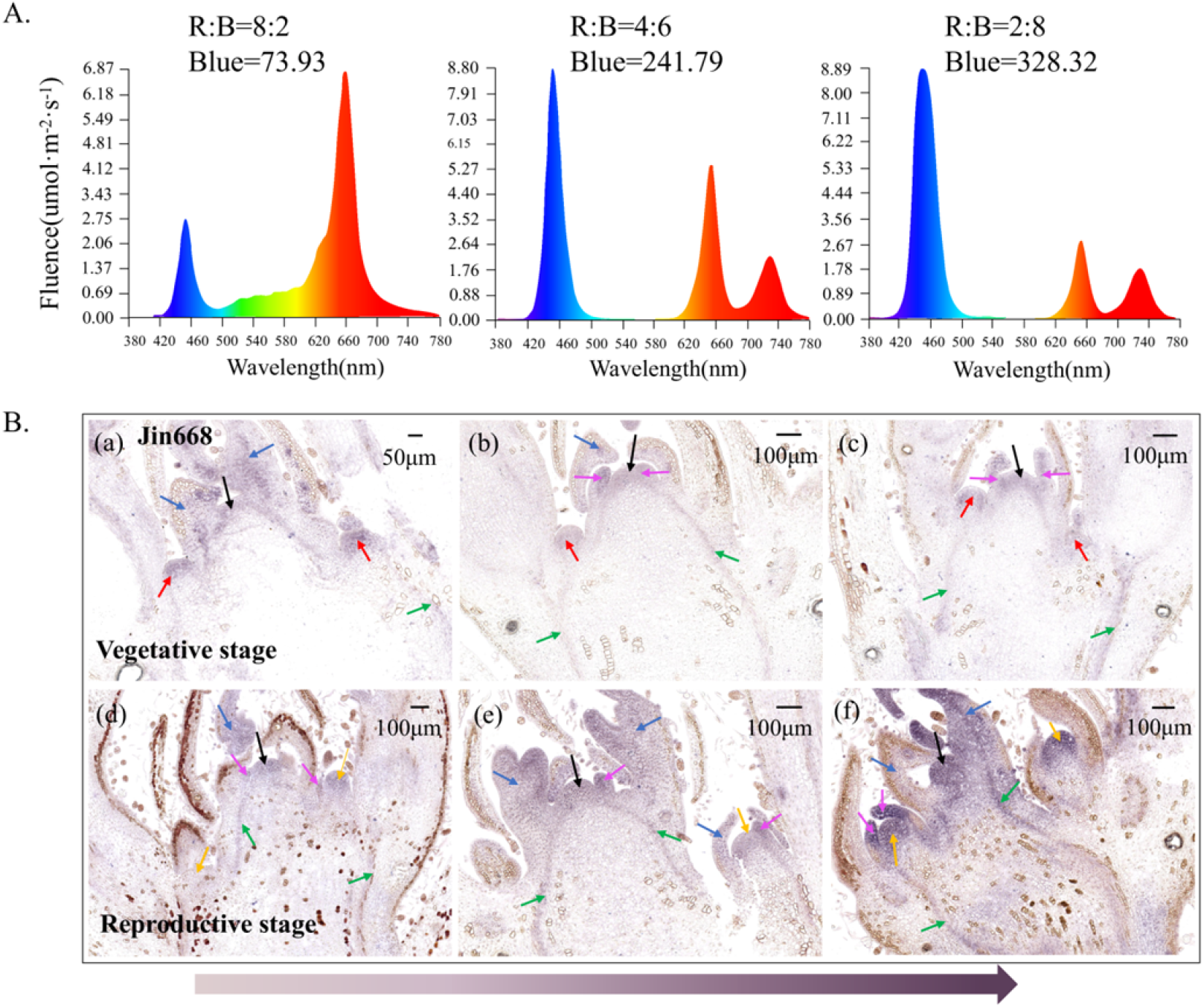
ISH was used to localize *GhCO* in the SAMs of cotton cultivar Jin668 under different R:B ratio treatments. **(A)** Light spectral compositions of different red:blue (R:B) ratio treatments. The R:B ratio and blue light fluence rate are shown above each light spectrum. **(B)** (a-c) Representative micrographs of longitudinal sections of SAMs for GhCO during the vegetative stage, with constitutive expression in the LP (the pink arrow), AM (the red arrow), FM (the orange arrow), SAM (the black arrow) and TL (the green arrow) under R:B = 2:8 (a), R:B = 4:6 (b) and R:B = 8:2 (c). (d-f) Representative micrographs of longitudinal sections of SAMs for GhCO during the reproductive stage under R:B = 2:8 (a), R:B = 4:6 (b) and R:B = 8:2 (c) treatments.

To further study the function of *GhCO* in cotton SAM stem cell fate decisions, *Ghco* mutants were obtained using the CRISPR–Cas9 system with two different sgRNAs (Fig. 9A-B), and mutations were tracked using the Hi-TOM platform (Fig. 9C). Importantly, sgRNA1 was located at the main functional region of the *CO* gene (B-box domain) (Fig. 9A). Phenotypic analysis showed a later flowering time but much more flowers in the mutant plants (Fig. 9D-E), suggesting that *GhCO* is involved in light signal functions in balancing flower primordium differentiation in quantity and timing.

**Fig. 9.**
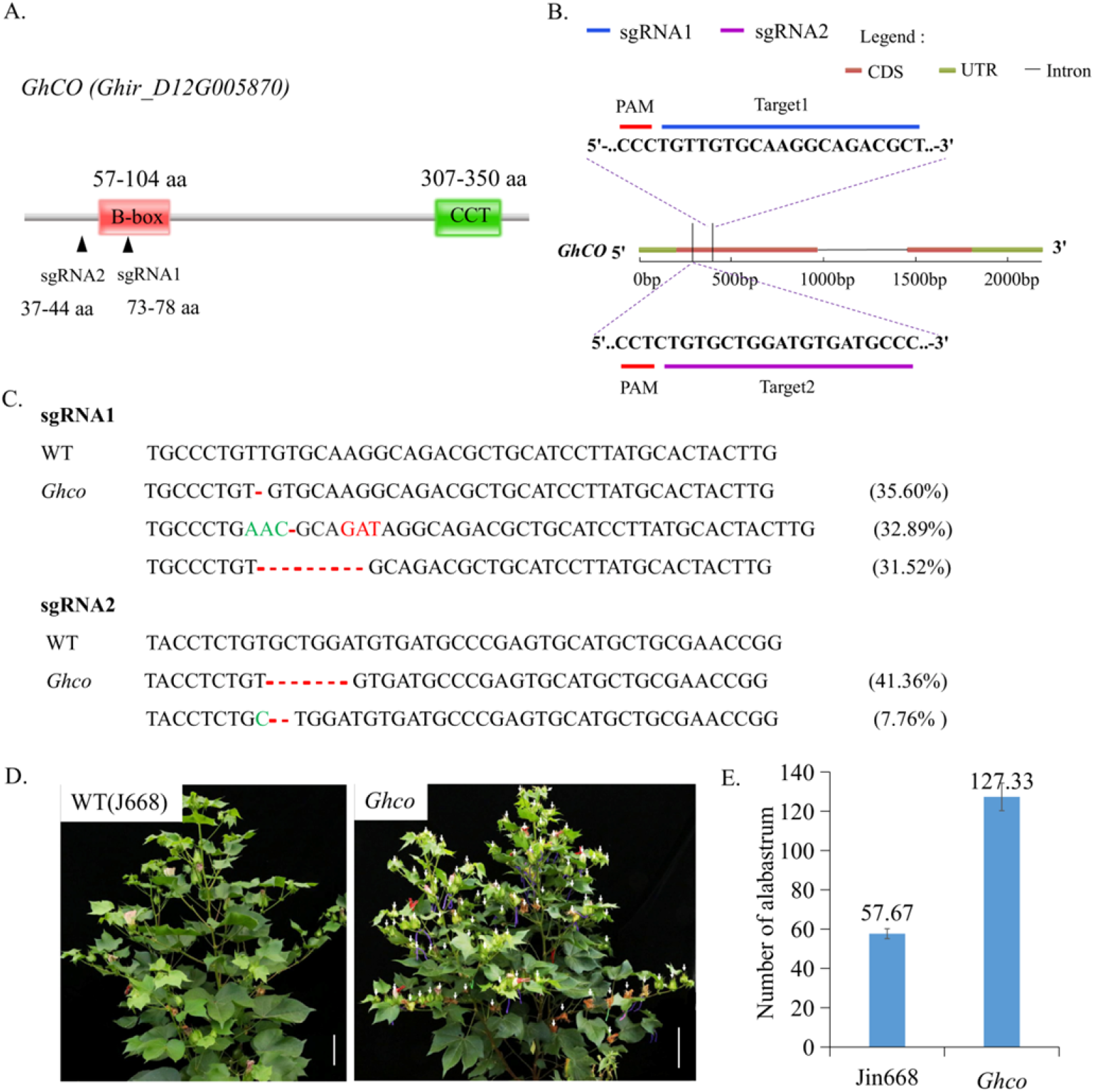
*Ghco* mutants created by the CRISPR–Cas9 strategy. **(A)** Schematic of the functional domains of *GhCO* predicted by SMART. The B-box (red box) and CCT domain (green box) are indicated. Black triangles, sgRNA1/2. **(B)** Schematic of the two small guide RNAs (sgRNAs, blue and purple bars) designed to target *GhCO* loci. PAMs are indicated by red bars. **(C)** Sequencing results showing indel mutations in the *GhCO* knockout plants. The percentages in parentheses represent gene editing efficiency on the right of sequences. **(D)** Phenotypes of the *Ghco* edited plant. White arrow, flowers or flower buds; scale bar, 10 cm. **(E)** The number of alabastrum statistics of Jin668 and *Ghco*. Data are represented as the mean ± SD (n = 3).

### Blue light signalling accelerates the transition from vegetative to reproductive development of cotton SAM

Previous studies have shown that CRYs, as blue light receptors in plants, participate in light inhibition of hypocotyl elongation and long-day promotion of floral initiation and photomorphogenesis (Ahmad and Cashmore, 1993; Guo et al., 1998; Franklin, 2016; Lyu et al., 2021). Moreover, WGCNA showed that the blue light receptor *GhCRY1* was co-expressed with *GhCO* and *GhFKF1* (Fig. 10A); therefore, *GhCRY1* may play a significant role in the response to light during initiation from the vegetative to reproductive stages of cotton SAMs. To confirm the function of blue light signals in the transformation from vegetative to reproductive development stages of cotton SAM, the treatment of the early-reproductive cultivar QS and late-reproductive cultivar XLZ, as well as CRISPR–Cas9 acceptor line YZ1 were treated with 400 μmol·m^−2^·s^−1^ total light fluence at 8:2, 4:6 and 2:8 R:B ratios and showed that a relatively high blue light proportion promoted flowering in all above three cotton cultivars (Fig. 10B). These results implying that blue light signalling accelerates the transition from vegetative to reproductive development of cotton SAM and facilitates flowering.

**Fig. 10.**
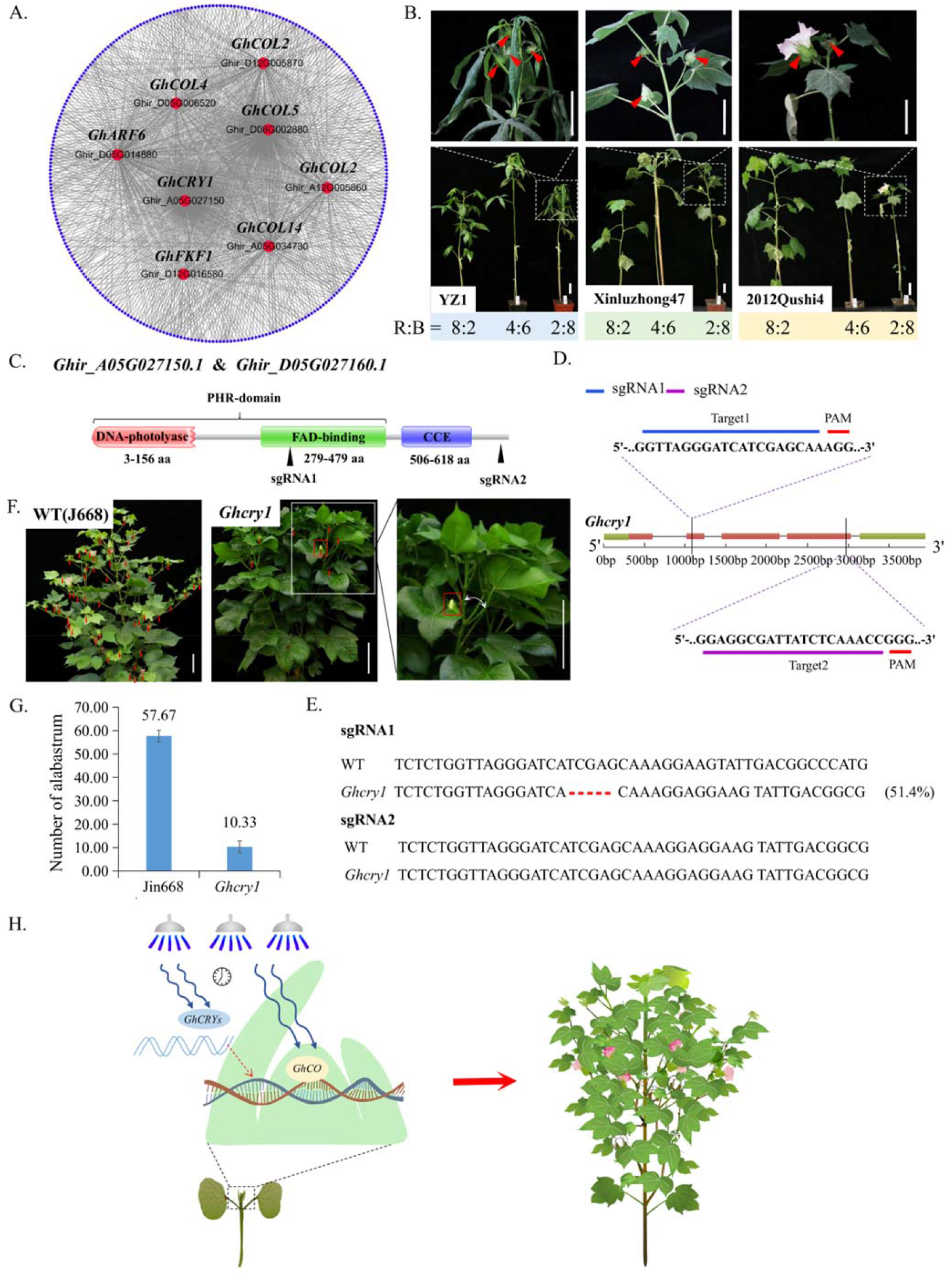
Blue light accelerates flowering of cotton. **(A)** WGCNA of all 4560 DEGs by Cytoscape showing a significant enrichment of known boundary-specific genes in response to flowering development. Red indicates reproductive development-related regulators defined according to the GO notion. Dots around the circle represent other co-expressed genes. Lines represent relationships. The white and black fonts represent the gene abbreviations and IDs, respectively. **(B)** Phenotypes of different cotton cultivars YZ1, XLZ and QS grown under LD conditions (16 h light/8 h dark) with different R:B ratios 2:8, 4:6, 8:2). Red arrows, flowers or flower buds; scale bar, 5 cm. **(C)** Schematic of the functional domains of *GhCRY1* predicted by SMART. The DNA-photolyase domain (red box), FAD-binding domain (green box) and CCE domain are indicated, and the DNA-photolyase and FAD-binding domains are also regarded as PHR domains. Black triangles, sgRNA1/2. **(D)** Schematic of the two small guide RNAs (sgRNAs, blue and purple bars) designed to target *GhCO* loci. PAMs are indicated by red bars. **(E)** Sequencing results showing indel mutations in the *Ghcry1* knockout plant. The percentages in parentheses represent gene editing efficiency on the right of sequences. **(F)** Phenotypes of the *Ghcry1* edited plant. Red boxes, flowers; two-way white arrows, angle between lateral branches and main stem; scale bar, 10 cm. **(G)** The number of alabastrum statistics of Jin668 and *Ghcry1*. Data are represented as the mean ± SD (n = 3). **(H)** Regulation models of the SAM differentiation mechanisms in response to blue light during cotton growth and development.

Furthermore, to confirm whether *GhCRY1* responds to light signalling, *GhCRY* (*Ghir_A05G027150* and *Ghir_D05G027160*) mutants were created by CRISPR–Cas9-mediated gene editing with two sgRNAs targeting their respective conserved functional domains (Fig. 10C-D). The mutations were identified by using the Hi-TOM platform (Fig. 10E). The knockout of *GhCRY1* in cotton not only led to a notable reduction in flowering number but also to a significant increase in vegetative branches along with a smaller bifurcation angle and flowering time delay (Fig. 10F-G). These results suggest that *GhCRY1* coordinates SAM stem cell differentiation by responding to light signalling, accelerates the floral primordium and increases the primordium number. Taken together, our results demonstrate that cotton SAM stem cells respond to blue light by *GhCRY1* and *GhCO* in coordinating the differentiation direction of cotton SAM stem cells between the vegetative primordium and reproductive primordium. Regulating the proportion of blue light in the growth environment or high *GhCO* accumulation in SAM stem cells can promote the transformation from vegetative to reproductive growth.

## Discussion

SAM stem cells orchestrate the balance between stem cell proliferation and organ initiation for postembryonic shoot growth, give rise to vegetative or reproductive primordia and reshape plant architecture (Jiang et al., 2013; Lu et al., 2013; Knauer et al., 2019). Many studies have been performed on the effect of the fate of apical stem differentiation regulated by hormones in various plants, such as rice, maize, tomato and grass (Wang and Li, 2011; Wang et al., 2012; Jiang et al., 2013; Zhang and Yuan, 2014). Light has also emerged as a key regulator of vegetative to reproductive transition (Turck et al., 2008; Song et al., 2015). To date, the cellular biological mechanism by which light signals regulate the differentiation fate of apical meristem cell clusters is not as complete as those in studies on hormones (Shani et al., 2006; Dong et al., 2013); these studies mainly focus on the influence of floral primordium initiation time (Zhang and Yuan, 2014). Cotton plant architecture and flowering time are determined and influenced by the fate of differentiation of the meristem and affect mechanized cotton production and yield. To this end, the illustration of light signal mechanisms on SAM differentiation has been urgently required for the improvement of cotton production efficiency.

In our study, we inspected the specific effects of light signals and key factors on cotton development and morphogenesis. The connection between light and cotton SAM differentiation was demonstrated, further elucidating the multilayer mechanisms contributing to cotton SAM fate decisions. Using the paraffin sectioning technique, dynamic changes in the cell morphology of cotton SAM during vegetative transformation into reproductive growth were observed, and the TFs *GhCO* and blue light receptors *GhFKF1* and *GhCRYs* were highlighted through transcriptome analysis. Moreover, we proposed an anatomically dynamic development module in regulating light signal-adjusted SAM differentiation in cotton (Fig. 10H).

RNA-seq analysis of the 35 libraries derived from seven different cotton varieties from five different developmental stages elucidated a link between light stimulus and cotton SAM differentiation. The initiation from vegetative to reproductive growth requires a morphological change (Zhang and Yuan, 2014). Our observations suggest that a model exists in which a light signal is required for cotton SAM fate decisions during the initiation from vegetative to reproductive growth. Through classification of the TFs based on their functions, the TFs *GhCO*-*like*, *GhRAV* and *GhTALE* were highlighted.

Moreover, a previously unknown role of *CONSTANS* in postembryonic and shoot system development was revealed by using the CRISPR–Cas9 edited system and ISH of cotton SAMs. The functional mechanisms of *CONSTANS* as a hub in the signal integration of photoperiodic flowering have been extensively illustrated in *Arabidopsis*, rice and wheat (Valverde et al., 2004; Song et al., 2015; Shim et al., 2017). In contrast, the influence of light signals on the fate of SAM is rarely studied, and the roles of *GhCO* in cotton remain largely unknown (Song et al., 2015). The phenotype of *Ghco* mutant plants demonstrates that *GhCO* affects SAM fate decisions during vegetative to reproductive growth by inhibiting the differentiation of the floral primordium number. In *Arabidopsis*, *CONSTANS* acts in the phloem to regulate a systemic signal that induces photoperiodic flowers (An et al., 2004; Gloria Serrano, 2009). It is worth noting that the ISH results, not only under LD conditions but also under different R:B ratio treatments, showed that *GhCO* accumulated at the SAM stem cells and vascular cambium of cotton, and showed that *GhCO* responded to blue light and accelerated the initiation from vegetative to reproductive growth of cotton SAMs, with a mechanism different from that in *Arabidopsis* (An et al., 2004; Turck et al., 2008).

Breeders proposed the ‘green revolution’ with a *semidwarf* mutation (*sd1*) identified in rice, and since then, rice and wheat yields have soared. Maize, tomato and legume varieties have also been shown to be thriving on the way to the ‘green revolution’ (Eveland et al., 2014; Zhang and Yuan, 2014; Lyu et al., 2021). Proper plant architecture and flowering time are the main goals of the ‘green revolution’, and SAM differentiation studies are primarily a target for idealizing plant agriculture and flowering time (Teo et al., 2014; Wang et al., 2018). Cotton is an important cash crop that displays a conventional long growth period, but growing early-maturity cotton is needed for the current wheat (rapeseed)-cotton cropping systems in China; therefore, breeding early-maturity cotton cultivars is urgent. Our results show that by regulating light signal-responding genes, *GhCRY1*, *GhFKF1* and *GhCO* may provide a strategy to shorten the cotton growth period for the current double cropping system, using limited land to support a daily enlarged population.

## Materials and Methods

### Plant materials and growth conditions

To acquire RNA-seq materials, three early-maturing cotton cultivars, CCRI58 (C), 2012Qushi4 (QS), and CCRI50 (CR), and four late-maturing cultivars, Ganmian47 (G), Xinluzhong47 (XLZ), Eguangmian (EG) and TM-1 (TM), were planted in the greenhouse of Huazhong Agricultural University in Wuhan (114 °E, 30 °N), Hubei Province, China (Fig. S1). The plants were cultivated under long-day (LD, 16 h light/8 h dark) conditions. Each variety was grown in seven 565×375×80 mm^3^ basins, with 40 plants per basin (Fig. S1C). Five stages of the SAM stem cell clusters were collected between 16:00 and 18:00 in the afternoon every two days (Song et al., 2015). For each stage, 15-18 SAM stem cell clusters of the cotton seedlings were pooled together and flash-frozen in liquid nitrogen and stored in a freezer at −80 °C (Fig. 1I).

To investigate the agronomic traits of the CRISPR–Cas9-edited mutants, the wild-type plants Jin668 and YZ1 and *GhFKF1*, *GhCO* and *GhCRY1* edited lines were grown in parallel with a row spacing of 55 cm and a plant spacing of 15 cm in the field of Huazhong Agricultural University.

### Tissue sectioning, staining, and imaging

The shoot apex was removed from the shoot tips, which were subsequently immersed in 50% FAA (50% absolute ethanol, 10% 37% formaldehyde solution, 5% acetic acid) and vacuum infiltrated three times for 15 min to fix the tissues. After infiltration, the solution was replaced with fresh FAA solution and postfixed at 4 °C for at least 12 h. Fixed samples were dehydrated with different ethanol concentrations (30, 50, 70, 95, and 100%) for 1 h. Overnight, tissues were embedded in 100% ethanol three times, 3/4 ethanol and 1/4 xylene, 1/2 ethanol and 1/2 xylene, 1/4 ethanol and 3/4 xylene, 100% xylene twice and then paraffin. The samples were successively immersed in xylene and 1/4 paraffin for at least 12 h, and then transferred to refined paraffin wax three times for 3 h. After embedding and cleaning, tissues were placed onto a microtome tissue holder, sectioned into 7 μm thick slices with a Thermo Scientific sliding microtome (Microm HM 340 E) and dried on a 37 °C heating plate for 2 h. The coverslips were mounted with resin and examined under a Zeiss Axio Scope A1 biological photomicroscope.

### RNA-seq and data analysis

For analysis of the mRNA profiles of SAM stem cells, seedlings were grown under LD conditions (16 h light/8 h dark) at 28-30 °C. Then, SAM stem cell clusters (1.5 mm in length) were collected and flash-frozen in liquid nitrogen. The SAM stem cells from 15-18 seedlings were pooled for each stage of each cultivar (Fig. 1I and Fig. S2C and D). Total RNA was captured with TRIzol (Invitrogen). Libraries were prepared using an Illumina TruSeq RNA Sample Prep kit following the manufacturer’s recommendations. The experiments were performed at Personalbio Gene Technology Co. Ltd. (Nanjing, China). After removing low-quality reads with Trimmomatic (Bolger et al., 2014), clean reads were mapped to the TM reference genome (Wang et al., 2019) by HISAT2 (Kim et al., 2019), and gene expression levels were calculated as fragments per kilobase per million (FPKM) by String Ties (Pertea et al., 2015). OmicShare tools, a free online platform for data analysis (www.omicshare.com/tools), was used to identify differentially expressed genes (DEGs) (P < 0.05 and log2 |fold-change| > 2) (Pertea et al., 2016).

Gene Ontology (GO) enrichment analysis of DEGs was determined using the OmicShare tools. The calculated *P* value was subjected to FDR correction, taking FDR ≤ 0.05 as a threshold. GO terms meeting this condition were defined as significantly enriched GO terms in DEGs. Moreover, Venn diagrams, heatmaps, Sankey diagrams and volcano plots were also generated by OmicShare tools.

### RNA isolation and RT-qPCR analysis

Total RNA was extracted with TRIzol (Invitrogen) reagent, and 3 μg of total RNA was used to generate complementary DNA using an Oligo (dT) 18 primer by TransScript® II One-Step gDNA Removal and cDNA Synthesis SuperMix (TransGen). Transcriptional levels of genes were determined using Premix Ex Taq (Takara) and analysed with a Quantstudio 6 Flex system. Two microlitres of 10-fold diluted cDNA was used as the template and amplified with TB Green® Premix Ex Taq (Takara) in a 20 μl volume quantitative PCR, which was pre-denatured at 95 °C for 30 s, followed by a 40-cycle program (95 °C 5 s, 60 °C 30 s, per cycle). Expression levels were normalized to *GhUBIQUITIN7* as an internal control to standardize RNA content. The primers used in this study are listed in Table S6 and S7.

### Analysis of protein and gene structures

The protein sequences were obtained from the Cotton FGD database. Then, the *GhFKF1*, *GhCO* and *GhCRY1* protein structures were visualized by using the pfam database. Similarly, the gene sequences were obtained from the Cotton FGS database. The gene structures were then visualized by using SMART online tools.

### Vector construction and genetic transformation

The gene editing strategy was performed using the CRISPR–Cas9 technique as previously described (Wang et al., 2018). Briefly, we selected fully developed seeds of the Jin668 and YZ1 cultivars and disinfected them with mercury at a concentration of 1 g mercury bichloride/1000 ml ddH_2_O for 10 min. The sterilized seeds were grown in a shading incubator for 3-5 days at 28 °C. Then, we cut the hypocotyl into sections as explants with a sharp scalpel. To construct the vectors, we searched for and identified 23 bp target sites (5’- N20NGG-3’) within exons of the *GhFKF1*, *GhCOR1* and *GhCO* genomic sequences according to the CRISPR-P online tool (Liu et al., 2017) and then evaluated each candidate site for target specificity on the website of potential off-target finder (http://crispr.hzau.edu.cn/CRISPR2/). We subcloned two independent sgRNAs targeting the target genes into the modified pRGEB32-GhU6.7-NPT II vector (Wang et al., 2018).

Then, the vectors were transferred into Agrobacterium (*Agrobacterium tumefaciens*) strain EHA105 by the electro transformation method. Explants were immersed in the EHA105 inoculum for 2-3 min with occasional agitation and then transferred to cocultivation plates for 48 h at 20 °C in shading conditions. Then, explants were cultured on callus initiation medium (2,4-D) with the explants laid flat on the medium under 12 h light/12 h dark photoperiod conditions at 27 °C and subcultured for 25 days in fresh medium. When the embryoids formed, they were transferred to plant induction medium. When seedlings developed at least one true leaf, they were transferred to rooting medium (Fig. 5A). The transgenic plants were then subjected to the Hi-TOM platform to evaluate whether mutations occurred (Liu et al., 2019). The sgRNA and HI-TOM primers used in this study are listed in Table S9 and S10.

### Co-expression network analysis of DEGs

To detect the groups of DEGs with similar expression patterns, a total of 4066 DEGs were used for weight gene co-expression network analysis (WGCNA). Co-expression modules were discovered using the WGCNA (v. 1.66) package in R software with default settings, except that the power was 12 and the minimum module size was 10 (Langfelder and Horvath, 2008). Subsequently, the topological overlap matrix (TOM) was calculated for hierarchical clustering analysis. Finally, a dynamic tree cut algorithm was implemented to identify gene co-expression modules. Then, the modules were visualized using Cytoscape 3.6.0 (Shannon et al., 2003).

## Accession numbers

Sequence data from this article can be found in the Cotton FGD database or the GenBank/EMBL libraries with following accession numbers: GhFKF1 (Ghir_D12G016580), GhCRY1 (Ghir_A05G027150), GhGI (Ghir_D02G006910) and GhCO (Ghir_D12G005870), GhCO-like (Ghir_A12G005860), GhNF-YB (Ghir_A07G020670), GhNF-YA (Ghir_A02G012520), GhCDF1 (Ghir_D12GO19090), GhHD-ZIP (Ghir_A12G022820), GhNAC (Ghir_D12G017690), GhRAV (Ghir_A12G018050), GhTALE (Ghir_D06G002490), GhC2H2 (Ghir_D04G012350), GhARF (Ghir_D05G014880), GhbHLH (Ghir_A11G003520), GhFAS2 (Ghir_A11G023990), GhCAT2 (Ghir_A11G025010), GhSVP (Ghir_A06G003180), GhAHP1 (Ghir_D13G019090), GhMCM3 (Ghir_D05G015210), GhGRP1A (Ghir_D02G022800), GhSOC1 (Ghir_A11G009010), GhEMB2247 (Ghir_D12G000040), GhGID1B (Ghir_D12G000700), GhELF3 (Ghir_D07G002430), GhPHOT2 (Ghir_D12G001220), GhMCCB (Ghir_A12G002340), GhELF4 (Ghir_A10G024010), GhVIP5 (Ghir_A10G002030), and At2g30600 (Ghir_D05G018350).

## Supplemental Data

**Supplemental Figure S1.** Growth stages of cotton for RNA sequencing.

**Supplemental Figure S2.** Heatmap of all differentially expressed genes (DEGs) that show correlated changes in gene expression clustered based on their expression trends.

**Supplemental Figure S3.** The transcription factors (TFs) analysis of DEGs.

**Supplemental Figure S4.** The photoperiodism pathway is required for initiation from vegetative to reproductive development.

**Supplemental Table S1.** The growth duration of cotton varieties with different maturities.

**Supplemental Table S2.** PCCs calculated of expressed genes from the At subgenomes.

**Supplemental Table S3.** PCCs calculated of expressed genes from the Dt subgenomes.

**Supplemental Table S4.** Functional classification of all 405 TFs of 4561 DEGs.

**Supplemental Table S5.** TFs involved in significantly enriched biological process GO terms, including flowering development, response to hormone catabolism, oxidation related, dimerization, cell differentiation, defence responses and ageing in selection sweeps for Sankey diagram analysis.

**Supplemental Table S6.** Primers for qRT–PCR of circadian rhythm-related genes.

**Supplemental Table S7.** Primers for qRT–PCR of photoperiod pathway-related genes.

**Supplemental Table S8.** The probe sequence of *in situ* hybridization (ISH).

**Supplemental Table S9.** sgRNA target sequences.

**Supplemental Table S10.** HI-TOM test primers for gene editing efficiency.

**Supplemental Table S11.** Statistics of the alabastrum numbers of WT Jin668 and YZ1 plants and transgenic *Ghgi*, *Ghfkf1* and *Ghco* plants.

**Supplemental Table S12.** The raw data in FPKM.

## Acknowledgements

We would particularly like to acknowledge Prof. Lili Tu from Huazhong Agriculture University for her valuable guidance regarding my research. We are also extremely grateful to the cotton team of the State Key Laboratory of Huazhong Agricultural University for their support and help in this study. This project was supported financially by funding from the National Key R&D Program of China (2020YFD1001004) and the China Agriculture Research System (Grant No. CARS-15-06).

